# AusAMF: database of arbuscular mycorrhizal fungal communities in Australia

**DOI:** 10.1101/2024.09.16.612857

**Authors:** Adam Frew, Jeff R. Powell, Meike K. Heuck, Felipe E. Albornoz, Christina Birnbaum, John D.W. Dearnaley, Eleonora Egidi, Luke Finn, Jarrod Kath, Kadri Koorem, Jane Oja, Maarja Öpik, Tanel Vahter, Martti Vasar, Stephanie Watts-Williams, Yuxiong Zheng, Carlos A. Aguilar-Trigueros

## Abstract

**Motivation:** Arbuscular mycorrhizal (AM) fungi are integral to plant nutrient acquisition, carbon cycling, and ecosystem resilience, yet our knowledge of their biogeography is severely limited, especially in the Southern Hemisphere. Australia, despite its landmass and unique geoecological characteristics, has been vastly undersampled, leaving a significant gap in our understanding of AM fungal diversity and distribution. The *AusAMF* database was created to address this deficiency, the first release comprises AM fungal community data from 610 sampling locations across mainland Australia and Tasmania, collected between 2011 and 2023. Using standardised sampling, DNA extraction, sequencing methods and platforms, this database provides a robust resource for exploring spatial patterns in AM fungal diversity, community composition, and the ecological drivers shaping AM fungal biogeography. The *AusAMF* database will continue to be updated and maintain standardised approaches to facilitate future research into plant-mycorrhizal interactions, nutrient cycling, and to understand the broader role of AM fungi in ecosystem processes. The data here will provide the foundation for more informed management and conservation efforts in Australia while providing valuable data for global-scale analyses.

**Main types of variables contained:** Georeferenced occurrence and abundance of high-throughput amplicon sequences of arbuscular mycorrhizal (AM) fungi.

**Spatial location and grain:** Australia. Decimal degrees between 0.000001 – 0.1 resolution.

**Time period and grain:** 2011-2023. Month and year of sampling.

**Major taxa and level of measurement:** Arbuscular mycorrhizal fungi identified to family, genus, and virtual taxon (VT). Geographic occurrence and amplicon sequence abundance.

**Software format:** Interact with data via online application. Dataset available as .csv files and raw sequencing data as .fastq files.

## 1. INTRODUCTION

The majority of terrestrial plants form symbiotic relationships with arbuscular mycorrhizal (AM) fungi (Smith & Read, 2008). In this relationship, AM fungi colonise the plant roots while simultaneously establishing extensive mycelial networks in the soil. As obligate symbionts, AM fungi rely on carbon provided by the host, which can account for as much as 17% of the total carbon fixed by certain host plants (Harris *et al*., 1985). The fungi facilitate access to essential soil nutrients, particularly phosphorus (P), a nutrient that commonly limits plant productivity. Phosphorus acquisition through AM fungi can supply up to 80% of the host’s P nutrition (Marschner & Dell, 1994). Beyond nutritional benefits, the symbiosis can also enhance plant resistance to insect herbivory and pathogens and improve tolerance to abiotic stress (Delavaux *et al*., 2017; Frew *et al*., 2022).

These fungi, recognised for their significant role in plant nutrition and survival, are also crucial in shaping plant community assembly, influencing primary productivity, and driving carbon and nutrient cycling. Given the morphological (Aguilar-Trigueros *et al*., 2019) and functional (Delavaux *et al*., 2017; Frew *et al*., 2022) diversity of AM fungi, it is essential to understand the abiotic and biotic factors that shape their biogeography. Such understanding is crucial for accurately predicting variations in their roles across ecosystems and for estimating large-scale patterns in global carbon cycling and primary productivity. Despite their ecological significance, knowledge of AM fungal biogeography remains limited in comparison to that of plants and other fungal taxa (Tedersoo *et al*., 2014).

Of the research that has been conducted to determine AM fungal biogeography, sampling efforts are strongly biased towards Europe, North America, and China, resulting in an uneven understanding of the biogeography of AM fungi. This disparity was made particularly evident in the release of the GlobalAMFungi database (Větrovský *et al*., 2023; release 1.0), which compiled sampling data across the globe from 100 studies focused on AM fungal composition. Vast regions of the Southern Hemisphere are evidently undersampled, with the soil AM fungal communities of Australia represented by only 32 samples, accounting for just 0.8% of the global soil samples collated. This makes Australia the most undersampled continent, excluding Antarctica.

This undersampling of the Australian continent represents a major gap in our understanding of AM biogeography. Australia comprises over 10% of the Southern Hemisphere’s landmass and, due to its Gondwanan origin, shares many soil and climate characteristics with large parts of Africa and South America (Flores-Moreno *et al*., 2023). However, Australia is also notable for harbouring the oldest continental crust and exhibiting an extraordinarily high level of species endemism, with approximately 93% of its plant species being endemic (Orians & Milewski, 2007). Many Australian plants are also capable of forming associations with more than one type of mycorrhiza (i.e., endomycorrhiza, ectomycorrhiza; Teste *et al*., 2020), while a significant diversity of plant taxa in Australia do not form associations with mycorrhizal fungi at all (Lambers *et al*., 2011). The ancient soils and unique biotic characteristics of Australia suggest that significant insights could be obtained from better quantification of AM fungi across its landscapes (Frew & Aguilar-Trigueros, 2023), yet it remains one of the least studied regions.

As with other fungi, molecular methods represent the state-of-the-art approach to characterise AM fungal diversity (Tedersoo *et al*., 2022). However, for most research, documenting the biogeographic patterns of AM fungi requires methods distinct from those employed for other major fungal groups (Větrovský *et al*., 2023). For the majority of fungi, research on their diversity can be adequately achieved using the internal transcribed spacer (ITS) region of the rRNA gene cluster, which is recognised as the most appropriate marker for investigating community composition and diversity in most fungal lineages through amplicon sequencing (DNA metabarcoding) (Schoch *et al*., 2012). However, AM fungi possess highly variable ITS copies, leading to probable overestimations of species richness and strong biases towards certain lineages in studies using ITS primers (Kohout *et al*., 2014; Öpik *et al*., 2014; Lekberg *et al*., 2018).

As such, the 18S small subunit (SSU) and the 28S large subunit (LSU) regions of the rRNA gene cluster are often the most appropriate markers for describing AM fungal diversity (Stockinger *et al*., 2010; Kohout *et al*., 2014). Although the debate regarding the choice of barcode markers remains ongoing (Delavaux *et al*., 2021; Tedersoo *et al*., 2022), most studies on AM fungal community diversity and composition have employed regions of the SSU gene. This region, being less variable than the ITS, provides conservative estimates of AM fungal richness, ensures robust recovery across AM fungal families, and facilitates analyses of phylogenetic relationships (Öpik *et al*., 2014; Thiéry *et al*., 2016). Additionally, the SSU is the primary marker utilised by Maarj*AM* (Öpik *et al*., 2010), a curated database of AM fungal SSU sequences, which links sequences to phylogenies and assigns them to reference ‘virtual taxon/taxa’ (VT), thereby enhancing comparability across different studies.

To fill the gap in our knowledge on biogeography of AM fungi, we present the first release of a continental-scale database of AM fungal communities based on SSU marker. The database derives from 610 sampling locations across mainland Australia and Tasmania (**Fig. 1**), collected between 2011 and 2023. The samples span all major climate zones in Australia (**Fig. 2**) making this dataset ideal to infer biogeographic patterns of these fungi. A key feature of the database is the use of a standardised protocol for processing all soil samples, enhancing the reliability of comparisons drawn from the data. All samples were collected from the top 10 cm of the soil layer, air-dried, and stored with silica to maintain dryness until processing. Each sample underwent the same DNA extraction procedures, followed by sequencing of amplicons of the SSU region using the primer set WANDA (Dumbrell *et al*., 2011) and AML2 (Lee *et al*., 2008), performed by Illumina NextSeq at the same sequencing facility. The raw sequencing data, in fastq format, are publicly available from the Sequence Read Archive (accession PRJNA1154677).

**Figure 1.**
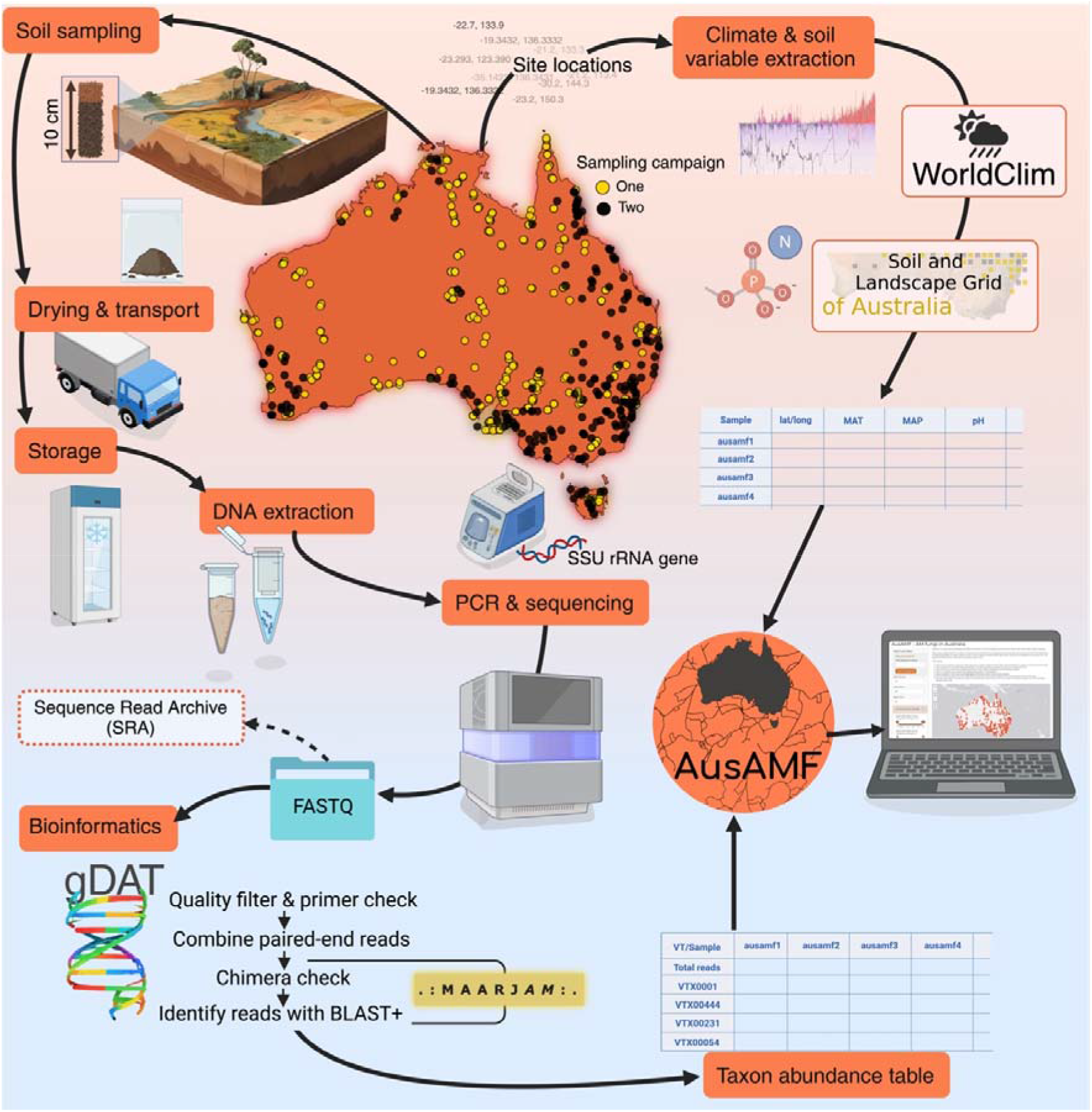
Workflow in generating data for *AusAMF*. Site locations recorded and soil samples taken from within the top 10 cm of soil. Soil is air-dried and stored with silica gel until transport to laboratory where samples are stored with silica gel in temperatures between 3-18. The DNA from soil is extracted and polymerase chain reactions used to amplify region of the small-subunit (SSU) using the WANDA and AML2 primer set (Lee *et al*., 2008; Dumbrell *et al*., 2011), after which the amplicons are sequenced using the Illumina NextSeq platform. Raw sequence data are openly available from the sequence read archive (SRA) of NCBI. Bioinformatic processing is performed using the gDAT pipeline (Vasar *et al*., 2021) and identification by referencing Maarj*AM* (Öpik *et al*., 2010). Climate and soil variables for each sample location are sourced from WorldClim and the Soil and Landscape Grid of Australia. These data are combined with AM fungal community sequencing data for each location and made available through the online *AusAMF* database (www.ausamf.com). Figure generated using biorender and freepik.

**Figure 2.**
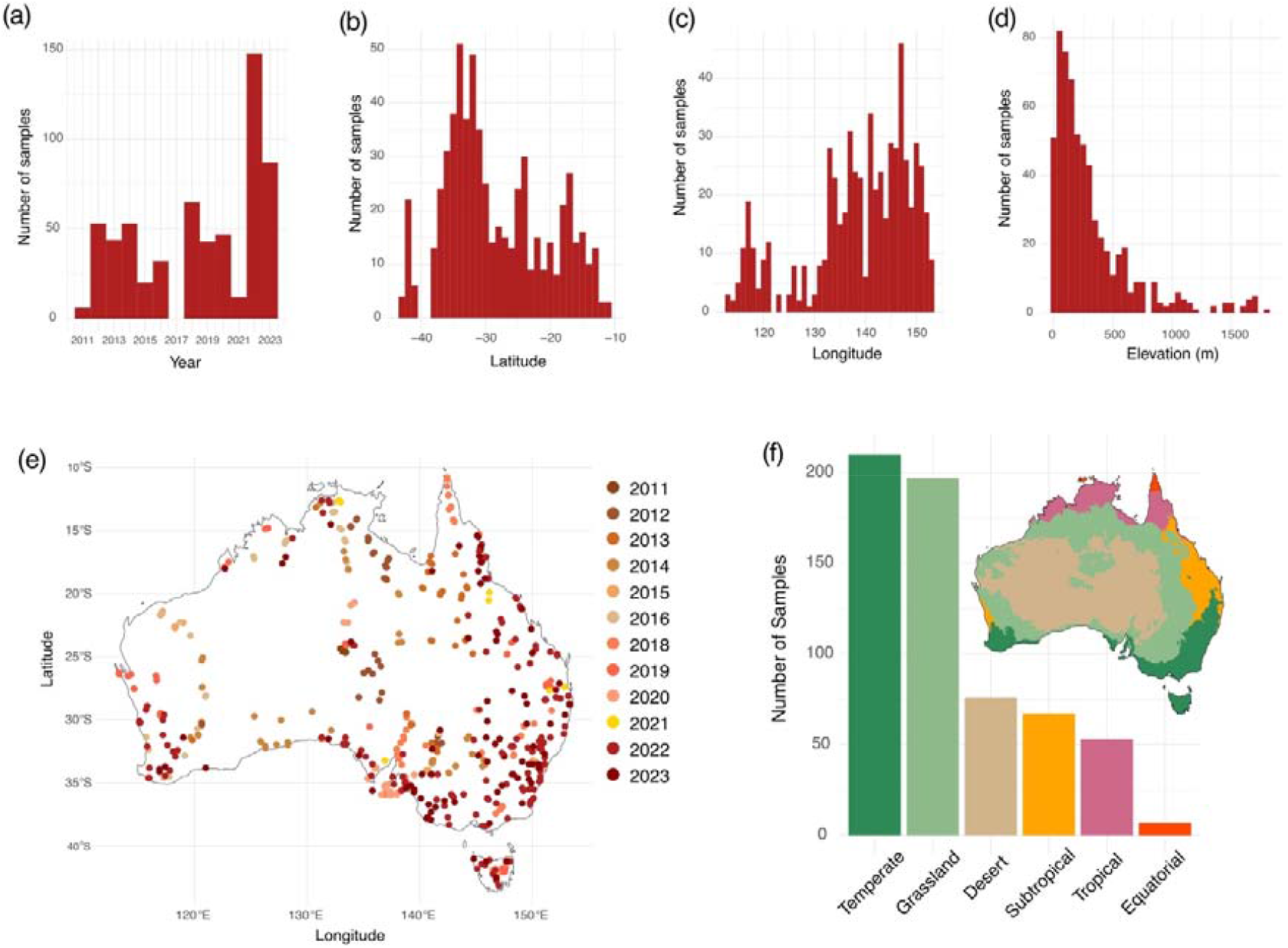
The distribution of samples in *AusAMF* by (**a**) year of sampling (**b**) latitude (**c**) longitude (**d**) elevation. (**e**) Map of sample locations in Australia coloured by year of sampling and (**f**) the number of samples from each of the major climate zones.

In addition, we processed these raw sequencing data using gDAT (Vasar *et al*., 2021), a bioinformatic pipeline specifically optimised for analysis of SSU amplicons from AM fungi, and taxonomy was assigned to VT (Öpik *et al*., 2010). For each sample, the spatial coordinates of sampling locations were used to download climate and soil variables from public repositories. These environmental data were combined with the AM fungal community data to allow users easy access to information that can be relevant for further analyses.

The *AusAMF* database will be updated with AM fungal community data from additional sites and locations when available. These additional data will remain as consistent as possible with the methods we describe below regarding soil sample collection, storage, DNA extraction, and sequencing. Users can also participate and provide soil samples or DNA, provided the sampling and DNA extraction procedures are comparable with those used here. Therefore, the database will provide spatial and temporal data on AM fungal communities across Australia, with continuous updates that will steadily expand both its depth and geographic scope.

## 2. METHODS

### 2.1 Soil sampling

The Australian arbuscular mycorrhizal fungal database (*AusAMF*) comprises high-throughput SSU sequence data from samples of soil collected across the Australian mainland and the island of Tasmania. Soil sampling for this initial dataset release (version 1.0) was conducted through two major sampling campaigns spanning from 2011 to 2023.

The first sampling campaign utilised sites from the Terrestrial Ecosystem Research Network (TERN) between 2011 and 2021, following the methods described in White *et al*. (2012). In brief, at each site, nine soil samples were collected from the top 3 cm of the soil profile within 100m x 100m plots across 375 locations in Australia included in this dataset. During sampling, gloves were worn, and instruments were cleaned with ethanol, methylated spirits, or 20% commercial bleach between samples to prevent contamination across samples and sites. Soil samples were air-dried and stored with silica gel in sealed bags or screw-top sample containers to maintain dryness. These samples were stored at a temperature-controlled facility (18 °C) until they were transported to Western Sydney University (Richmond, NSW, Australia) in 2023, where they were stored with silica at 4 °C until DNA extraction.

The second sampling campaign, conducted over 12 months from August 2022 to September 2023, involved soil collection from 235 sites, spanning both agricultural and non-agricultural environments across Australia. At each site, six soil sub-samples were collected from the top 10 cm of the soil within a 25 m² plot. These were collected at equal distances along a transect, or if not possible, samples were collected equally spaced in a grid from the 25 m^2^ plot. Samples were collected wearing gloves, litter and debris were removed from the surface prior to taking the soil. Any equipment and instruments used were cleaned with ethanol, methylated spirits, or at least 20% commercial bleach, prior to sampling. The soil samples were pooled and air-dried at room temperature (avoiding direct sunlight and temperatures above 39 °C) and sealed in plastic bags with silica gel before being transported to Western Sydney University (Richmond, NSW, Australia), where they were stored at 4 °C until DNA extraction.

### 2.2 Environmental variables

Site locations as latitude and longitude were recorded. Those samples collected from sampling campaign one (TERN) provide locations to 4 decimal places, soil collections from sampling campaign two were provided by third parties comprising various researchers, land managers, and volunteers. The longitude and latitude for these locations are provided to a lower resolution (0.1 decimal point). Environmental variables included in the dataset are mean annual temperature (MAT), mean annual precipitation (MAP), elevation, soil total nitrogen, soil total phosphorus, soil pH (CaCl_2_), soil organic carbon (SOC), and soil available phosphorus. The MAT, MAP, and elevation data were retrieved from the WorldClim Bioclimatic variables (Fick & Hijmans, 2017), which provide environmental data based on the average for the years 1970 – 2000.

The soil variables were retrieved from the Soil and Landscape Grid of Australia, an open-access database using digital soil mapping and extensive soil sampling across the continent (Terrestrial Ecosystem Research Network, 2024). Details of the sampling, analyses and mapping methodologies are available for total nitrogen (Rossel *et al*., 2014), total phosphorus (Malone & Searle, 2024), pH (Malone & Searle, 2021), soil organic carbon (Wadoux *et al*., 2022), and available (Colwell) phosphorus (Zund, 2022).

### 2.3 DNA extraction and sequencing

The DNA from each soil sample was extracted from 250 mg of soil using DNeasy Powersoil Pro Kits (Qiagen, GmBH, Hilden, Germany) according to the manufacturer’s instructions, apart from the final step where DNA was eluted using UltraPure™ Distilled Water (Invitrogen, MA, USA). Sequencing was performed by the Ramaciotti Centre for Genomics (UNSW Sydney, Australia). DNA underwent amplification by polymerase chain reaction (PCR) of the small-subunit (SSU) ribosomal RNA gene using the WANDA (5′-CAGCCGCGGTAATTCCAGCT-3′; Dumbrell *et al*., 2011) and AML2 (5′-GAACCCAAACACTTTGGTTTCC-3′; Lee *et al*., 2008) primer set with Illumina overhang sequences. For this, thermal cycling conditions for PCR were as follows: 95 °C for 2 min; 27 cycles of 95 °C for 60 s, 54 °C for 60 s, and 72 °C for 60s; with a final extension of 72 °C for 10 min. DNA quality control gels were used to confirm successful amplification. All samples were purified using the Agencourt AMPure XP Beads (Beckman Coulter, CA, USA). The PCR products were then amplified in an indexing PCR reaction with Nextera compatible unique dual indexes (Integrated DNA Technologies, Coralville, IA, USA). The thermal cycling conditions were: 95 °C for 3 min; 8 cycles of 95 °C for 30 s, 55 °C for 30 s, and 72 °C for 30s; with a final extension of 72 °C for 5 min. Quality control gels were again used to confirm successful amplification and all samples were normalised using SequalPrep Normalization Plate Kit A1051001 (Thermo Fisher Scientific, MA, USA). All samples were pooled with an equal volume and cleaned using the Agencourt AMPure XP Beads (Beckman Coulter, CA, USA). The cleaned sample pool was quantified by Qubit quantification assay (Thermo Fisher Scientific, MA, USA) before being sequencing on an Illumina NextSeq 1000 platform using the Illumina P1 600 XL kit.

### 2.4 Bioinformatics

Bioinformatic data analysis and processing was conducted using the graphical downstream analysis tool (gDAT) for analysing rDNA sequences (Vasar *et al*., 2021). The default settings of the gDAT pipeline are optimised for the analysis of AM fungal SSU sequences. We followed the defaults as outlined in Vasar *et al*. (2021). The raw reads (88,022,138) were retained if they carried the correct primer sequences (WANDA and AML2; allowing one mismatch for each). Quality filtering was done by checking the average base quality against a threshold (at least 30) combined with a sliding window average quality check to trim low-quality regions, leaving 71,464,445 cleaned reads. Paired-end reads were combined using *FLASH* (Magoč & Salzberg, 2011) with default minimum 10-bp overlap and 75% overlap identity threshold, resulting in 69,447,662 (97.18%) combined pairs. Although gDAT provides the option, we did not pre-cluster reads into operational taxonomic units (OTUs). Chimera checking was conducted with *VSEARCH* (Rognes *et al*., 2016) in denovo and reference mode which checked each sequence individually against the reference database, in this case, Maarj*AM* (Öpik *et al*., 2010). This identified 163,094 chimeras (0.2%), 69,258,958 non-chimeras (99.7%) and 25,610 borderline sequences. The AM fungal sequences, and their taxonomies, were identified using BLAST+ (Camacho *et al*., 2009) where each individual read was identified in comparison to the Maarj*AM* database (Öpik *et al*., 2010) as a reference using the nucleotide local pairwise alignment *BLASTN* algorithm. This identified sequences to phylogenetically defined taxonomic units called virtual taxa (VT). A taxon abundance table was then generated at 97% identity matching and 95% alignment.

### 2.5 Data exploration and visualisation

All data exploration and analyses were conducted using R version 4.3.3 (R Core Team, 2024).

The MAP, MAT, elevation, soil pH (CaCl_2_), soil organic carbon, total nitrogen, total phosphorus, and available phosphorus were extracted for each sample location using the ‘raster’ (Hijmans, 2023) and ‘sp’ (Bivand & Gomez-Rubio, 2013) packages in R. Frequency histograms, scatterplots, mean and standard error plots were generated using ‘ggplot2’ (Wickham, 2016). Maps of Australia were produced using the ‘rnaturalearth’ package (Massicotte & South, 2023). Rarefaction curves to compare sequencing depth to number of VT were made using *rarecurve* from ‘vegan’ (Oksanen *et al*., 2015). Plots showing proportional abundance of AM fungal families were generated using the *transform_sample_counts* and *psmelt* functions from ‘phyloseq’ (McMurdie & Holmes, 2013) together with ‘ggplot2’ (Wickham, 2016). In addition to observed number of VT, extrapolated AM fungal richness and Shannon diversity metrics were calculated (Hill numbers of order 0 and 1) using the ‘iNEXT’ package (Hsieh *et al*., 2016), to estimate diversity at complete coverage (Chao *et al*., 2014). The phylogenetic diversity was calculated (Faith’s phylogenetic diversity; Faith, 1992), which measures the sum total phylogenetic distance of each community. Species richness is typically highly correlated with raw phylogenetic diversity, therefore to help exclude the effects of richness the standardised effect sizes (SES) were calculated for phylogenetic diversity by way of *ses.pd* function from the ‘picante’ package (Kembel *et al*., 2010). The SES also allows the detection of non-random patterns of phylogenetic diversity.

The online interactive application for *AusAMF* was created using the ‘shiny’ package in R (Chang *et al*., 2024), together with packages ‘shinyBS’ (Bailey, 2022), ‘bslib’ (Sievert *et al*., 2024), ‘leaflet’ (Cheng *et al*., 2024), ‘DT’ (Xie *et al*., 2024), ‘dplyr’ (Wickham *et al*., 2023), ‘ape’ (Paradis & Schliep, 2019), ‘Biostrings’ (Pagès *et al*., 2024), ‘ggtree’(Yu *et al*., 2017), ‘readr’ (Wickham *et al*., 2024) and ‘ggplot2’ (Wickham, 2016).

## 3. Data description

### 3.1 Sample coverage

*AusAMF* release 1.0 includes high-throughput amplicon sequencing data from 610 sites in Australia sampled between 2011 and 2023 (**Fig. 2a,e**). Geographic coverage was slightly higher in the higher absolute latitudes (i.e. southern latitudes) and higher longitudes (**Fig. 2b,c**). The majority of sample sites sit < 500 m in elevation, which reflects most of the Australian continent (87% sits below 500 m). Site locations were classified to a climate zone based on a modified Köppen-Geiger classification system which incorporates information on the climatic limits of native vegetation, as the best expression of climate in an area (Stern *et al*., 2000). Temperate and grassland climate zones had the most sample coverage in the dataset (**Fig. 2f**).

### 3.2 Amplicon sequencing, diversity and composition of AM fungi in Australia

After quality filtering, primer and chimera checking, a total of 69,258,958 amplicon reads were obtained for further analysis. The mean and median number of sequences per sample was 113,539.3 and 73,975.5, respectively (**Fig. 3**). The individual sequences were identified to VT referencing the Maarj*AM* database (Öpik *et al*., 2010) which identified 2,336,123 VT reads across the samples from 610 sites.

**Figure 3.**
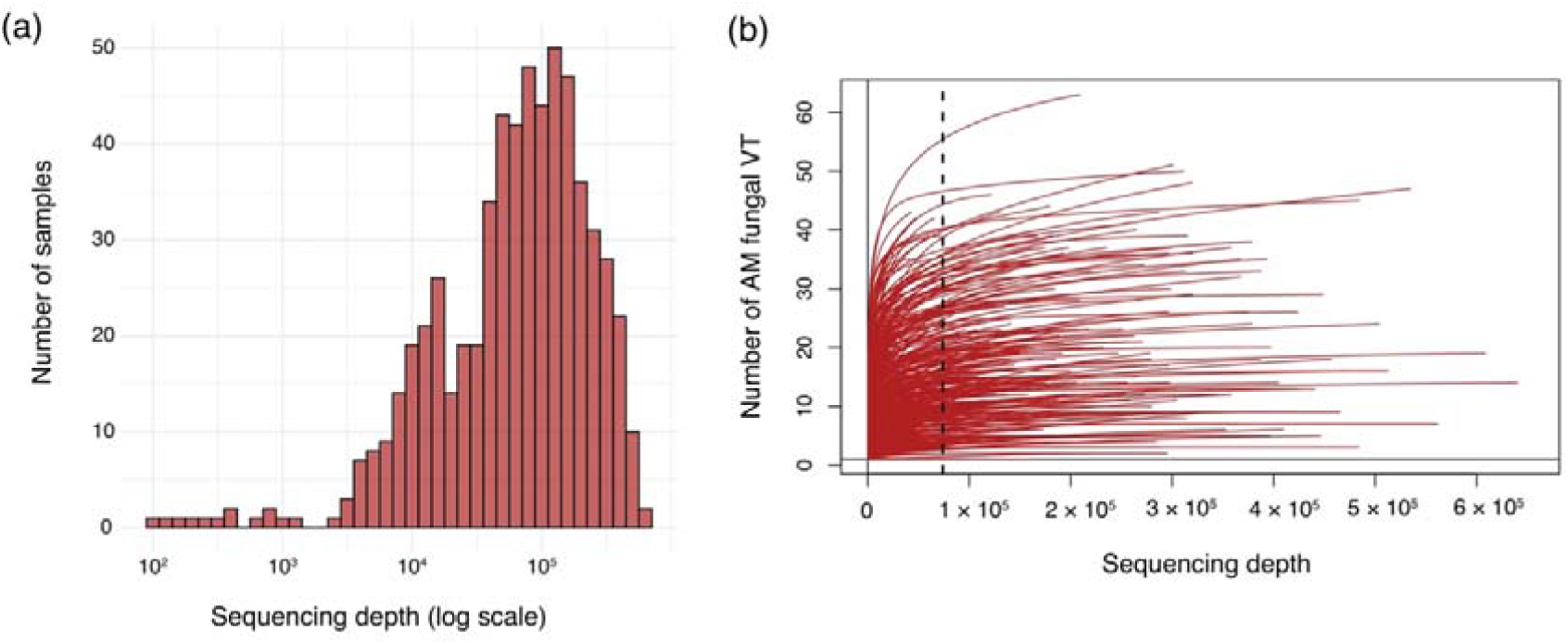
(**a**) The number of samples by sequencing depth and (**b**) the rarefaction curves for all samples showing accumulative number of arbuscular mycorrhizal (AM) fungal virtual taxa (VT) by sequencing depth.

In our dataset we identified 200 AM fungal VT across all samples, comprising nine families. Rarefaction curves comparing the number of sequences per sample to number of VT exhibited plateauing in most samples, suggesting an adequate sampling depth in most instances to capture AM fungal VT diversity. We have not rarefied data here, however researchers should carefully consider if and how the uneven sampling depth among samples might affect their research question(s), and how they might deal with this when conducting their own analyses (Tedersoo *et al*., 2022).

The mean VT richness (**Fig. 4d**) and extrapolated richness (**Fig. 4e**) exhibited similar patterns among climate zones. The mean VT richness was highest in tropical (22.64 ± 1.87; mean ± SE) and equatorial samples (21.14 ± 3.96; mean ± SE), while the lowest mean richness was in temperate samples (12 ± 0.68; mean ± SE). As richness does not account for the frequencies of VT, it can overweigh rare taxa in the community. Shannon index, in this case the extrapolated Shannon (Hill *q* = 1), takes frequencies into account but does not disproportionately favour rare or abundant taxa. Extrapolated Shannon (**Fig. 4f**) was also highest in tropical samples (6.56 ± 0.46; mean ± SE), however equatorial samples exhibited the lowest mean Shannon index (3.86 ± 1.16; mean ± SE). Distinct from the patterns in richness and Shannon index, the mean SES of phylogenetic diversity (**Fig. 4g**) was lowest in samples from equatorial (−1.11 ± 0.43; mean ± SE) and tropical climate zones (−0.12 ± 0.18; mean ± SE), and was highest in samples taken from the temperate region (0.96 ± 0.07; mean ± SE). In terms of compositional relative abundance (**Fig. 5**), the majority of sequences were from the family Glomeraceae (58.97%), while Pacisporaceae represented the lowest relative abundance of sequences (0.03%).

**Figure 4.**
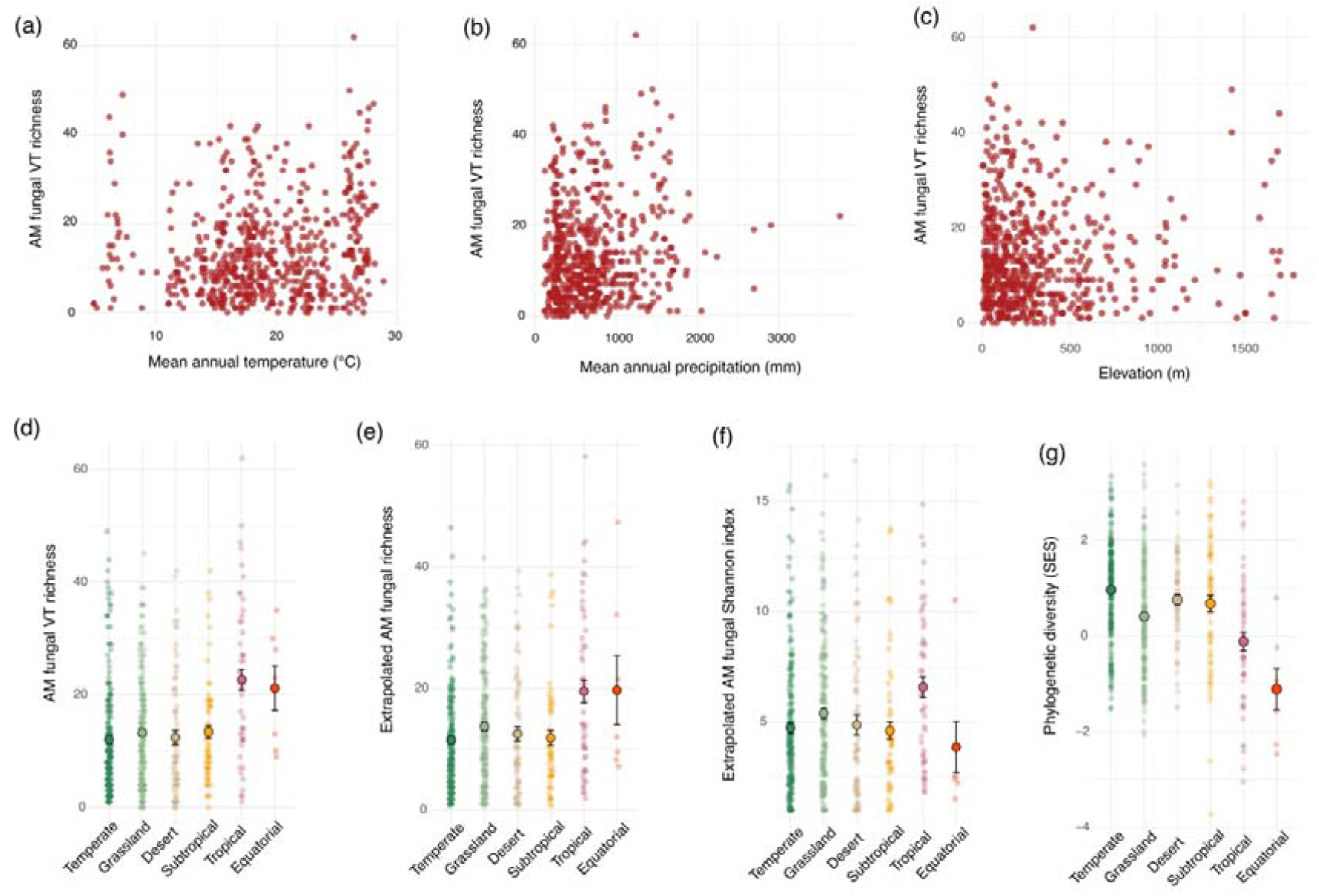
Scatterplots showing relationships between arbuscular mycorrhizal (AM) fungal virtual taxon (VT) richness and (**a**) mean annual temperature (**b**) mean annual precipitation, and (**c**) elevation. The (**d**) observed AM fungal richness, (**e**) extrapolated AM fungal richness (Hill number q = 0), (**f**) extrapolated AM fungal Shannon index (Hill number q = 1), and the (**g**) standardised effect sizes (SES) of phylogenetic diversity (Faiths) for AM fungal communities from the different major climate zone based on the modified Köppen-Geiger classification. Solid points and error bars in represent mean ± standard error which are overlaid on top of the raw data points.

**Figure 5.**
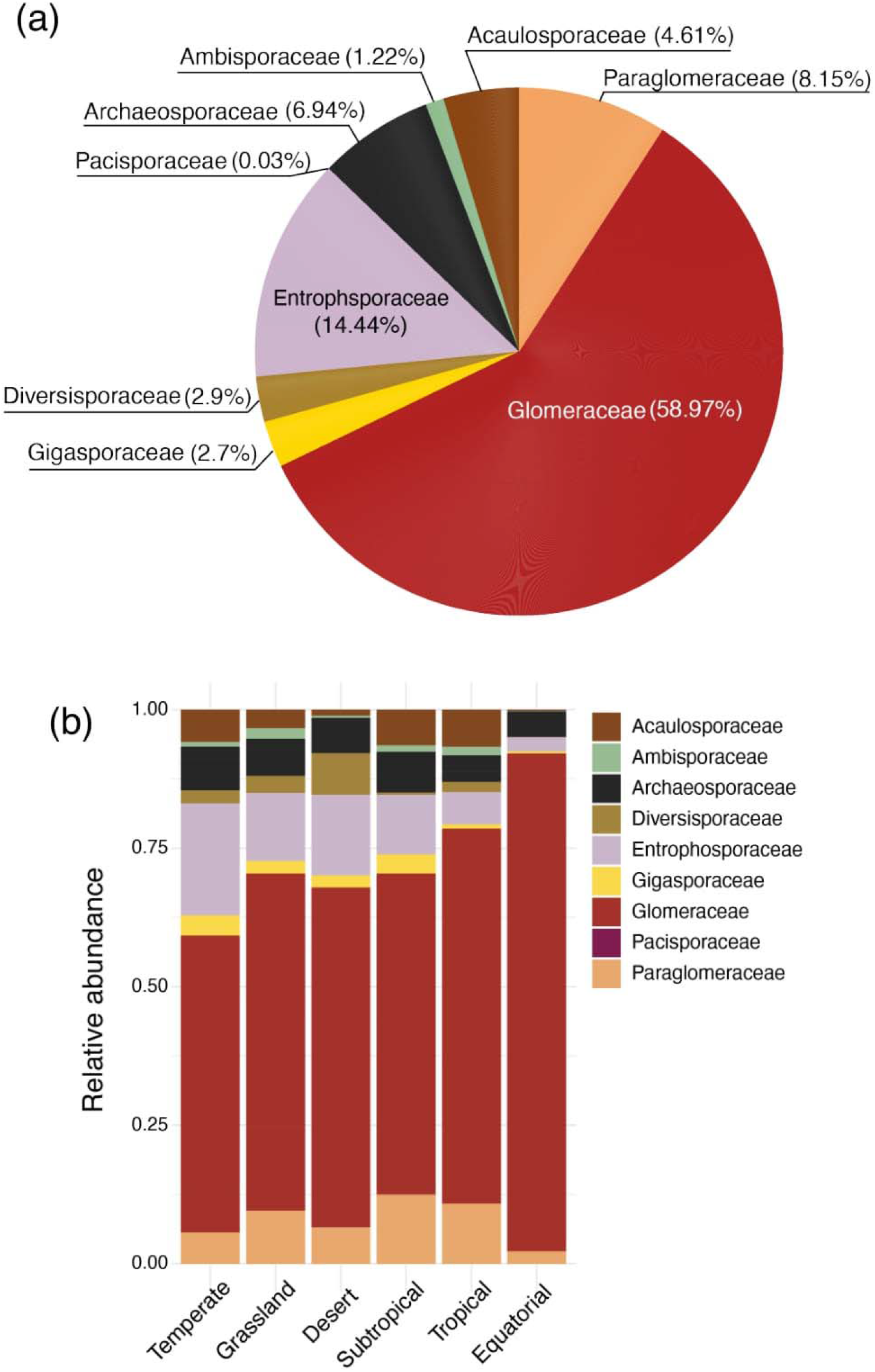
Relative abundance of arbuscular mycorrhizal (AM) fungal families within (**a**) *AusAMF* database and (**b**) the relative abundance of AM fungal families in the major climate zones based on modified Köppen-Geiger classification.

### 3.2 Database access

*AusAMF* can be accessed via the graphical user interface of the online application available at https://www.ausamf.com/, which is compatible with most browsers. From this platform, users can download a CSV file containing processed sequencing data for each sample along with associated environmental variables (Table 1). The application also provides options for visualising and filtering samples and AM fungal taxa, which can then be exported as a CSV file. For instance, users may visualise and download data for samples containing specific taxa (at the family, genus, or VT level) or filter samples based on environmental variables such as mean annual temperature (MAT), mean annual precipitation (MAP), or soil pH. Summary data are generated within the user interface when samples are selected, either by clicking on individual samples or by selecting samples on the map using the polygon tool. A phylogenetic tree is also generated for the AM fungal VT present in the selected sample(s). Additionally, the platform offers a search function that allows users to input a sequence, returning a table of representative VT sequences from the *AusAMF* dataset ranked by their similarity to the input sequence. The raw sequencing data are available for download from the NCBI Sequence Read Archive under accession PRJNA1154677, provided as paired fastq files for each of the 610 samples included in this first release.

**Table 1.** Summary of information included in the *AusAMF* database.

| Heading | Description |
| --- | --- |
| sample_number | Sample designation for the *AusAMF* database. These all start with ausamf followed by a numerical designation ### (e.g., ausamf015, ausamf524) |
| FASTQ_name | Sample name on the FASTQ files associated with the sample |
| latitude | The latitude of the location where the sample was taken |
| longitude | The longitude of the location where the sample was taken |
| year | The year the soil sample was collected |
| month | The month the soil sample was collected |
| koppen | The major climate zone classification, based on modified Köppen-Geiger classification system |
| MAT | Mean annual temperature (°C), calculated average between the years 1970-2000, for the location the sample was taken |
| MAP | Mean annual precipitation (mm), calculated average between the years 1970-2000, for the location the sample was taken |
| elevation | The elevation (m) for the location the sample was taken |
| total_N | Total nitrogen (%) in soil of the location the sample was taken |
| total_P | Total phosphorus (%) in soil of the location the sample was taken |
| avail_P | Available (Colwell) phosphorus (mg/kg) in soil of the location the sample was taken |
| SOC | Soil organic carbon (%) in soil of the location the sample was taken |
| pH_cacl | Soil pH (CaCl<sub>2</sub>) of the location the sample was taken |
| sampling_campaign | The sampling campaign the soil sample was collected from. The first version of *AusAMF* comprises two campaigns (see methods). This column is populated with either '1' (the first sampling campaign) or '2' (the second campaign) |
| cleaned_reads | The total number of sequence reads in the sample after quality filtering and chimera checking |
| total_abundance | The number of reads of the associated AM fungal VT in the sample |
| VT_ID | The virtual taxa (VT) identifier for AM fungi has the prefix 'VTX' followed by five numbers (e.g., VTX00142, VTX00105) |
| Family | Taxonomic assignment to family of the AM fungal VT |
| Genus | Taxonomic assignment to genus of the AM fungal VT |

## Acknowledgements

The authors thank TERN and the SLGA for providing access to soil sample collection, the Ramaciotti Centre for Genomics and S.P.U.N. The majority of this work was supported by an Australian Research Council (ARC) Discovery Early Career Researcher Award (DECRA; DE220100479) awarded to A.F. and also supported M.K.H. through a postgraduate research scholarship. J.R.P. was supported by an ARC Future Fellowship (FT190100590), S.J.W-W was supported by an ARC DECRA (DE210100908), as was E.E. (DE210101822). C.A.A-T was supported by an Academy Research Fellowship (21000058691) from the Research Council of Finland.

